# Detection, communication, and individual identification with deep audio embeddings: A case study with North Atlantic right whales

**DOI:** 10.1101/2025.07.11.664307

**Authors:** Irina Tolkova, Holger Klinck, Dana A. Cusano, Anke Kügler, Susan E. Parks

## Abstract

Anthropogenic noise has increased ambient sound levels across the globe, both underwater and on land. Among its many negative impacts, heightened noise can impair communication in vocal animals through acoustic masking. Conceptually, noise reduces the animal’s *communication space* – the area in which an individual animal can effectively convey information to a conspecific listener. Previous studies have estimated the communication space using sound propagation models and/or behavioral studies. However, studies frequently equate signal recognition with signal detection – a necessary but not sufficient precondition – thereby persistently overestimating spatial coverage and underestimating anthropogenic impacts. Measuring communication is inherently difficult, and varies with taxa, call type, and context, leading to significant data gaps in key parameters. We propose that deep learning creates an opportunity to estimate biologically-relevant communication, even for data-limited species. In particular, we present a case study with the critically endangered North Atlantic right whale (*Eubalaena glacialis*; hereafter NARW). Prior research has demonstrated that the upcall – a low-frequency contact call produced across ages and sexes – encodes individual identity. We therefore consider a dataset of NARW vocalizations recorded with on-animal archival tags, spanning 234 samples across 11 individuals from 3 sites. First, we demonstrate that audio embeddings from the BirdNET model can robustly distinguish individual right whales. Then, we simulate the effect of varying ambient noise levels to estimate signal excess for both signal detection and individual identification. Altogether, we hope this work provides both a methodological advance for individual identification and a framework for better understanding anthropogenic impacts on vocal wildlife.

## I. INTRODUCTION

Since the industrial revolution, anthropogenic noise has vastly reshaped global soundscapes. On land, transportation noise from automobiles and airplanes has given rise to elevated noise levels even in remote natural areas^1^. The oceans have experienced even greater change, due to exponential expansion of commercial shipping, oil and gas exploration, military sonar, underwater construction, and other sources^2,3^. In humans, chronic noise exposure has been associated with adverse health impacts ranging from increased risk of cardiovascular disease^4,5^ to cognitive impairment^6,7^. In animals, studies have documented extensive physiological and behavioral responses to noise^8,9^. Alongside effects on physiological health, increased ambient noise can mask vocalizations, impacting the ability of animals to communicate. Consequently, changes in vocal activity have been observed across taxa in response to increased noise, ranging from increasing call amplitude (known as the Lombard effect)^10–12^, increasing call frequency (thereby reducing masking by low-frequency noise sources)^13,14^, or changing call rate and duration^15,16^.

By obscuring a signal with noise, acoustic masking decreases an animal’s ability to communicate. Conceptually, masking reduces the animal’s “communication space” – the space in which the animal can effectively perceive and convey vocal information. Early studies aimed to quantify this region by measuring behavioral changes in response to call playback at different signal-to-noise ratios (SNR)^17–19^. While these studies yield numerical thresholds for a visible response, they fall short of truly measuring communication. Is the meaning behind the vocalizations still transmitted at the specified SNR thresholds, or are the animals simply detecting familiar sounds, too distant and faint to be understood?

Lohr et al^20^ shed some light on this question through a laboratory study of budgerigars and zebra finches trained to indicate both detection and discrimination of conspecific and hetero-specific calls. The authors demonstrated that call discrimination was indeed more challenging than detection, requiring over 3 dB higher SNR on average. Integrated with a simple model of acoustic propagation, this difference would imply a discrimination distance under 70% of the detection distance, and consequently a 2-dimensional discrimination space about half the size of the detection space. Building on these findings, Dooling et al^21^ suggested a conceptual framework with further tiered levels of communication: (1) on the outside, the detection space, in which an animal can detect a vocalization; (2) within it, the discrimination space, in which an animal can discriminate between multiple vocalizations; (3) next, the recognition space, in which an animal can recognize and interpret the vocalizations; (4) and finally, the innermost space of comfortable communication. In a basic propagation model assuming simple spherical spreading, these spaces can be imagined as concentric circles around the speaker or listener.

While proposed for birds, this framework can be translated across taxa, and has been considered for another group deeply impacted by anthropogenic noise: marine mammals^22^. Sound is the principal form of communication underwater – critical for social interaction, navigation, hunting and foraging, and other aspects of life. In the United States, the Marine Mammal Protection Act, enacted in 1972, mandates the assessment and mitigation of adverse impacts on marine mammals – aptly including the impacts of acoustic masking^23^. Accordingly, significant study has been devoted to establishing scientifically-grounded species-specific thresholds at which noise affects communication^22^. However, properly defining communication is a challenging and highly context-dependent question. Behavioral studies as performed with laboratory birds may be too effort-intensive and prohibitively invasive, if not unethical, to conduct with marine mammals. Indeed, a large-scale review of masking in marine mammals highlights “Studies on signal excess required for signal detection versus discrimination, recognition and comfortable communication” as a research need, though noting: *The signal excess needed for comfortable communication is difficult (if not impossible) to measure in animals*^22^.

Computationally, estimates of detection spaces are approached by formulating a sonar equation to calculate received sound levels, with appropriate terms for source levels, noise levels, and transmission loss, and modeling acoustic propagation to evaluate received SNR across an environment^22,24^. These studies recognize the need for an additional term to represent not just detection but information transfer; yet due to the lack of data, this term is either guessed or simply dropped. In consequence, the terms “active space”, “communication space”, and “detection space” are inconsistently defined in scientific literature, and frequently used interchangeably. This distinction is not just a conceptual concern, but carries practical consequences for conservation. As detecting a signal is a necessary but not sufficient condition for effective communication, studies that conflate the two will persistently over-estimate the communication space, and thereby under-estimate the adverse effects of anthropogenic noise in impact assessments.

In this work, we propose that machine learning can aid in bridging this gap. In the last decade, the field of bioacoustics has undergone a technological revolution from the widespread adoption of machine learning tools^25,26^. In particular, deep neural networks have enabled automated detection and classification of diverse animal vocalizations within long audio recordings, greatly expanding the spatial and temporal scales of acoustic analysis. Models trained on bioacoustic data have been shown to effectively generalize across taxa, from birds to marine mammals, and tasks, from species classification to one-shot search^27^. In particular, these models have shown very high sensitivity to acoustic features across scales, identifying characteristics of species, call types, regional dialects, and even individuals^28,29^. In this light, machine learning could also provide a new opportunity to quantify information transfer within animal vocalizations, filling a key gap in the study of communication.

Specifically, we present a case study for a Critically Endangered species known to be gravely impacted by anthropogenic noise – the North Atlantic right whale (*Eubalaena glacialis*; hereafter NARW). NARW are slow-moving, surface-dwelling baleen whales that live along the North Atlantic coast. An easy target for harpoons, NARW were hunted to near-extinction by the start of the 20th century^30^. While NARW received protections from commercial whaling in 1935, their population struggled to recover, increasingly faced with different lethal threats: collisions with ships and entanglement in fishing gear^31^. Now numbering fewer than 400 individuals, NARW are a crucial focus in marine conservation^32^.

While ship strike and entanglement are evident through scars, dragged gear, or necropsies, the effect of anthropogenic noise is understood to be severe but less visually apparent. As all whales, NARW rely on sound for communication, vocalizing in a vocabulary of low-frequency (*<* 500 Hz) calls^33^. Historically, this frequency range would have minimal interference from other sound sources, allowing these calls to propagate for many kilometers^34^. However, since 1950, commercial shipping has grown rapidly across the globe, with a ten-fold increase in gross tonnage by 2010^2^. Correspondingly, measurements of ambient noise in the 25-50 Hz band have increased by approximately 3.3 dB per decade^2,35–37^.

NARW have been shown to increase both amplitude and frequency of their calls in response to noise, which could reduce the effect of masking, but likely with an energetic cost^38^. Unsurprisingly, high noise levels have been linked to increased stress hormone production in NARW^39^, affecting individual welfare and likelihood of reproduction. The impacts of acoustic masking have wide-ranging impacts on a social species. Of principal concern are the impacts on mother-calf pairs; while raising young, mothers communicate with lower-amplitude calls, likely to avoid drawing attention to their vulnerable calves^40^, but thereby increasing the risk of losing each other amidst the noise^41^. Of the NARW call types, the most common call is the upcall: an upsweep from 50 to 350 Hz, 1-2 seconds in duration, that serves as a contact call for NARW of all ages and sexes^42^. While considered to be heavily steryotyped, an analysis of upcalls recorded on on-animal archival tags has shown that these vocalizations contain sufficient variability to distinguish individual whales with high accuracy^43^. Recordings of upcalls therefore provide a unique opportunity to quantify the transfer of one form of informational content – individual identity – and approximate the communication space of calling NARW under different noise regimes.

Overall, in this work, we aim to utilize deep learning to quantify communication and estimate thresholds for detection and recognition. This work has two key objectives. First, we build on prior work to demonstrate that individual NARW can be distinguished through upcalls in an automated framework, without the need for manual feature selection. Second, we show that the ability to distinguish individuals declines as background noise levels increase, quantify this relationship as a function of the signal-to-noise ratio (SNR), and compare the sound levels required for detection and individual identification. Methodologically, we address a collection of challenges: leveraging deep learning for a small dataset, verifying robustness to noise, and adapting to lower-frequency data. We then contextualize these results within prior estimates, and discuss the implications of our analysis for improving estimates of anthropogenic impacts on a critically endangered species.

## II. DATA ACQUISITION

We leveraged a dataset of acoustic recordings from three key regions of North Atlantic right whale range^44^: the Bay of Fundy, Canada^45–47^; Cape Cod Bay, Massachusetts, USA^48^; and the Southeastern US^49^. The data was collected with suction-cup-attached digital archival tags, including DTAGS^47^ and Acousonde tags^50^. For further detail on data collection, please reference McCordic et al (2016)^43^. We considered individuals with at least 10 upcall samples to avoid misleading accuracies over small sample sizes. Each upcall was resampled to a 2kHz sampling rate, normalized by the mean amplitude, bandpass filtered to a range of 50 to 500 Hz with a Butterworth filter, and standardized to a length of 3 seconds by zero-padding. In total, we considered 234 upcalls across 11 individuals.

## III. METHODS

Our methodology entails a series of analytical steps: calculation of embeddings, evaluation of individual identifiability, robustness to noise, modification of frequencies, and controlled variation of signal-to-noise ratio (SNR). First, to construct a meaningful lower-dimensional representation of NARW upcalls, we use audio embeddings from the pre-trained deep neural network BirdNET (described in Section III A). Our metrics for evaluating classification performance are described in Section III B. To ensure that noise is not a decisive confounding factor in our analysis, we apply de-noising, and analyze a noise dataset in parallel with the upcall dataset (Section III C). To optimize the performance of BirdNET for low-frequency whale calls, we apply and evaluate two forms of frequency modification (Section III D). Lastly, after demonstrating the potential for individual identification, we vary SNR to quantify the impact of ambient noise levels on the accuracy of both detection and idenification (Section III E).

### A. BirdNET Embeddings

A prior study by McCordic et al (2016) demonstrated that upcall characteristics encode individual identity along with age class^43^. In particular, the authors identified both frequency- and time-related discriminating features, such as the start and mean values of the fundamental frequency and duration. However, manually selecting acoustic metrics imposes assumptions and limits the patterns on the acoustic characteristics that can distinguish individuals. In this work, we investigate whether a fully automated, deep-learning-based approach can differentiate upcalls across individuals. While deep learning has revolutionized bioacoustic classification, sample sizes collected from tagging studies have typically been too small for training deep neural networks from scratch. However, networks trained on large, broad-scale datasets may be successfully applied to new, previously unseen categories and tasks through transfer learning^51^. One form of transfer learning proposes that a neural network trained for an audio classification task will internally represent meaningful acoustic features: deeper layers will extract increasingly refined characteristics, converted into an output classification label by the final layer(s). In this framework, the activations of a penultimate layer of a neural network would be meaningful “feature vectors” specialized for the dataset. These feature vectors, commonly termed *embeddings*, can then be visualized, compared, or analyzed further with simple classification algorithms.

For applications in bioacoustics, BirdNET is an openly accessible deep learning model and research tool for species-level classification of bird vocalizations, supporting over 6000 species globally^52^. Recent studies have demonstrated that BirdNET embeddings allow for high-performance transfer learning across taxonomically diverse vocalizations, including classification of bats, marine mammals, and frogs, as well as for diverse tasks, such as classification of dialects^27,28^. In fact, BirdNET embeddings have yielded higher classification accuracies for downstream bioacoustic tasks than more generic audio models, suggesting that the audio characteristics learned for bird song and calls are also meaningful for other animal vocalizations^27^. For this work, we therefore utilized BirdNET embeddings to serve as feature vectors. The embeddings were calculated using the command-line interface for BirdNET Analyzer v1.5.1, yielding vectors of 1024 features for each audio sample.

### B. Evaluation Metrics

After calculating BirdNET embeddings for all up-call samples, we evaluated the potential for differentiating individual right whales with these acoustic features. First, to visualize the distribution of the embeddings, we applied UMAP^53^ to reduce feature dimensionality from 1024 to 2, and visualized clustering structure within the distributions of embedded points. If embedded points associated with an individual clustered together, we could infer that tag-specific audio characteristics were a significant driver of the variation in upcall samples. Next, we evaluated the performance of a simple classifier – specifically, a linear support vector machine (SVM) – to distinguish individuals in the lower-dimensional embedding space. The performance of the SVM was compared with a baseline random classifier, which always predicted the most common class observed in the training data. As the low sample size rendered the division of the dataset into separate training and test datasets impractical, we instead evaluated leave-one-out cross-validation (CV) accuracy, consistent with prior work^43^.

### C. Denoising

One common pitfall in the analysis of data with deep learning methods is that models may learn and characterize aspects of a recording other than the desired target signal. As the recordings for an individual NARW were associated with a single tag in a particular environment, the upcalls were likely to contain individual-specific ambient noise profiles, creating the possibility that the BirdNET embeddings were learning characteristics of tag-specific noise rather than of individual-specific upcalls. To evaluate this effect, we constructed a “control” dataset of noise samples which was analyzed in parallel to the upcall dataset. Specifically, we selected 1-sec samples immediately before each upcall to represent the associated ambient noise profiles, and applied the above classification pipeline this additional dataset. If classification performance was driven primarily by upcall characteristics, we would expect the analysis of noise data to yield little to no clustering and low cross-validation accuracies.

Indeed, preliminary analysis showed high classification accuracies across both the upcall and noise datasets. We accounted for this complication by applying rigorous denoising to both datasets. Specifically, we calculated spectrograms for both the upcall and noise samples with a Hann window of length 256 samples (0.128 seconds) with 50% overlap, and extracted the rows representing the 50-500 Hz range. To account for the spectral characteristics of the ambient noise, we subtracted the time-averaged spectrum of the noise spectrogram from the magnitudes of each column of the associated call spectrogram. Similarly, to address impulsive or narrowband noises, we subtracted the mean magnitudes across both rows and columns. Lastly, time-frequency bins with a magnitude below a chosen threshold were set to 0; a range of thresholds was tested to select an optimal value. As the upcall is characterized by a narrow contour sweeping upwards in frequency, we anticipated for it to be robust to these subtraction steps. Finally, the phases of all time-frequency bins were set to the original values to preserve signal coherence, and the denoised spectrogram was mapped back to the time domain with an inverse Short-Time Fourier Transform. This denoising pipeline was applied to both the primary dataset of upcalls and the “control” dataset of ambient noise samples.

Then, to ensure that the classification results were representing characteristics of upcalls rather than of tag-specific noise, the analysis procedure was performed in parallel for the upcall samples and noise samples across a range of de-noising thresholds. At the threshold for which the CV accuracy of the noise samples was near-random, we could conclude that upcall accuracy was truly indicative of individual identification.

### D. Frequency Shifting

BirdNET embeddings were constructed as the model learned species-specific differences in bird calls and song. In consequence, these features could be expected to best represent the avian frequency range, and to likely have poorer sensitivity for acoustic features at low (*<* 500 Hz) frequencies. To account for this effect, we applied and compared two techniques for increasing the frequency of the upcalls prior to calculating embeddings. First, we considered a simple compression of the signal, shortening the signal in time and proportionately scaling the frequency range. We considered scaling factors from 2 (yielding a duration of 1.5 sec and frequency range of 100-1000 Hz) to 20 (yielding a duration of 0.15 sec and frequency range of 1000-10000 Hz) in increments of 2. Additionally, we considered increasing the frequency by 1 kHz while keeping duration constant through complex modulation. Specifically, we calculated a new signal *y* with values given by:

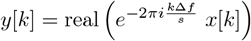

for all indices *k* = 1, …, *N*, where Δ*f* is the frequency shift to apply to the signal, *s* is the sampling rate, and *N* is the number of samples in the signal *x*. We again considered a range of shifts, from Δ*f* = 1 kHz to Δ*f* = 10 kHz in increments of 1 kHz. Lastly, a Butterworth high-pass filter with a cutoff of Δ*f* was applied to *y*[*k*] to prevent aliasing.

### E. Quantifying Detection and Recognition

After evaluating the ability to identify individuals across denoising parameters and frequency modifications, we aimed to quantify the effect of ambient noise levels on both detectability and individual recognition. First, we selected de-noised upcall samples for the best-performing denoising threshold and frequency modification, for which classification accuracy was quantified. Next, we added randomly-sampled white noise to achieve SNR values ranging from −30 to 30 dB at increments of 5 dB. Specifically, SNR was defined as the ratio of root-mean-square signal amplitude to noise amplitudes within the upcall band (50-500 or equivalent after frequency modification). For each SNR level, we repeated the above classification pipeline, calculating embeddings and quantifying CV scores. To quantify detectability, we evaluated binary classification, distinguishing upcall samples from noise samples, relative to a random accuracy of 50%. To quantify recognition, we evaluated individual classification, relative to a random accuracy of about 23% (the proportion of the largest class in the dataset). For robustness across random sampling, the generation of white noise was repeated ten times, providing a distribution of cross-validation scores for each SNR level.

## IV. RESULTS

### A. Effects of Denoising

The removal of ambient noise described in Section III C is demonstrated in Figure 1. In this example, this pre-processing approach effectively removed the overall ambient noise profile and persistent impulsive noise visible in the upcall recording. No harmonics above 500 Hz are visible due to a bandpass filter. By a threshold of 0.15, the noise sample becomes entirely collapsed to zero, while key aspects of the call sample are preserved.

**FIG. 1.**
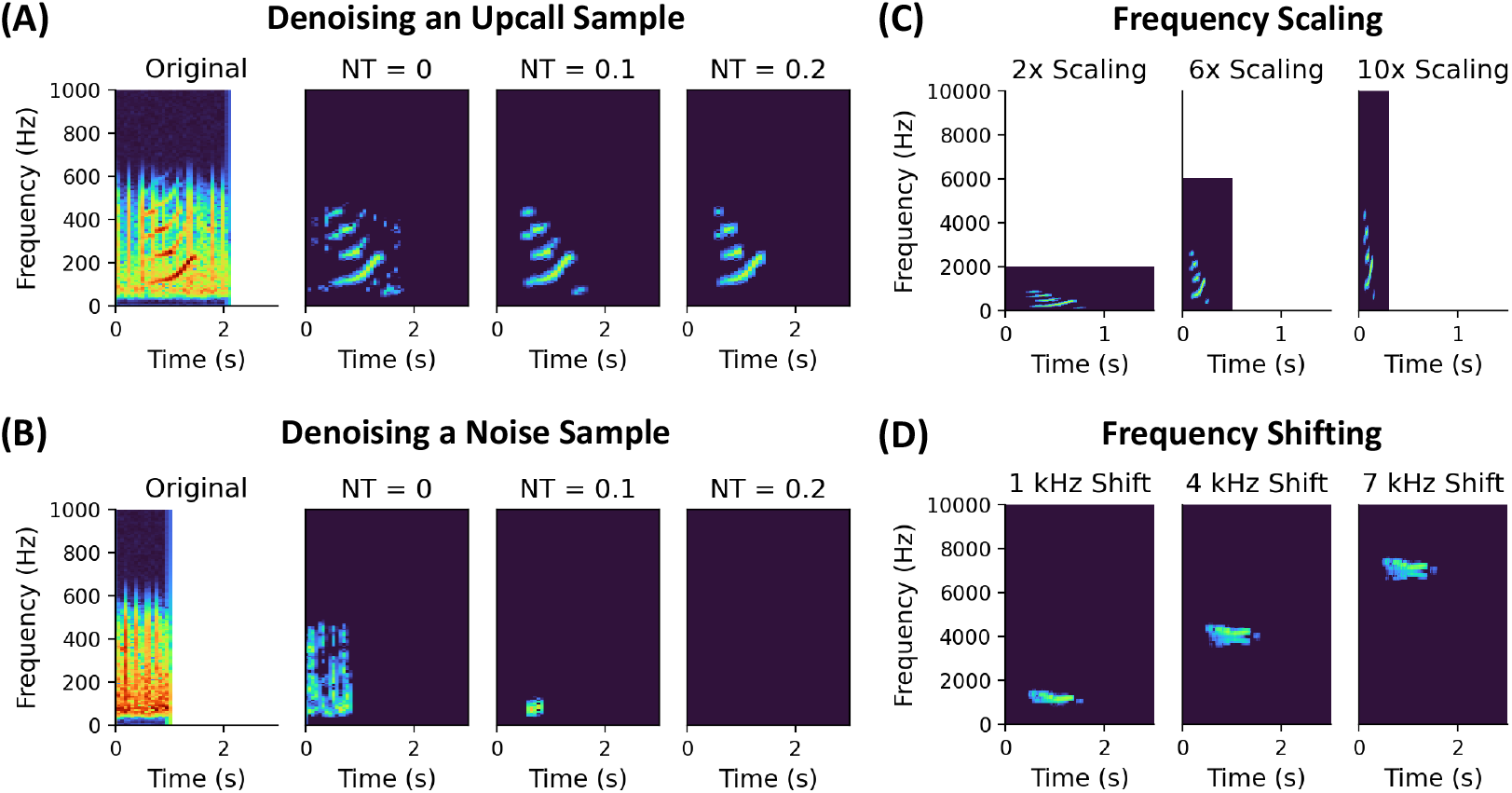
**(A)** The leftmost spectrogram displays an upcall sample with significant background noise. The following spectrograms display the sample after processing at different de-noising thresholds NT: 0, 0.1, and 0.2. **(B)** To de-couple classification of individual identity from classification of tag-specific noise, the same processing pipeline is applied to a corresponding noise sample for each upcall. **(C)** Frequency scaling with factors of 2x, 5x, and 10x is applied to a de-noised upcall sample. This approach compresses the duration and proportionally stretches the frequency range of the signal. **(D)** Frequency shifting at shifts of 1 kHz, 4 kHz, and 7 kHz is applied to a de-noised upcall sample. This approach maintains both the duration and difference between the maximum and minimum frequencies of the signal.

### B. Effects of Frequency Shift

Visualizations of frequency modification for an example upcall, both by speeding up and shifting up, are displayed in Figure 1. Correspondigly, the leave-one-out CV accuracies across denoising thresholds for both forms of frequency modification are displayed in Figure 2. Broadly, we find that accuracy can be improved by up to 10% through frequency modification. Overall, speeding up the calls was more effective than shifting up the calls, with an optimal speedup factor of 10x and an optimal shift of 2 kHz.

**FIG. 2.**
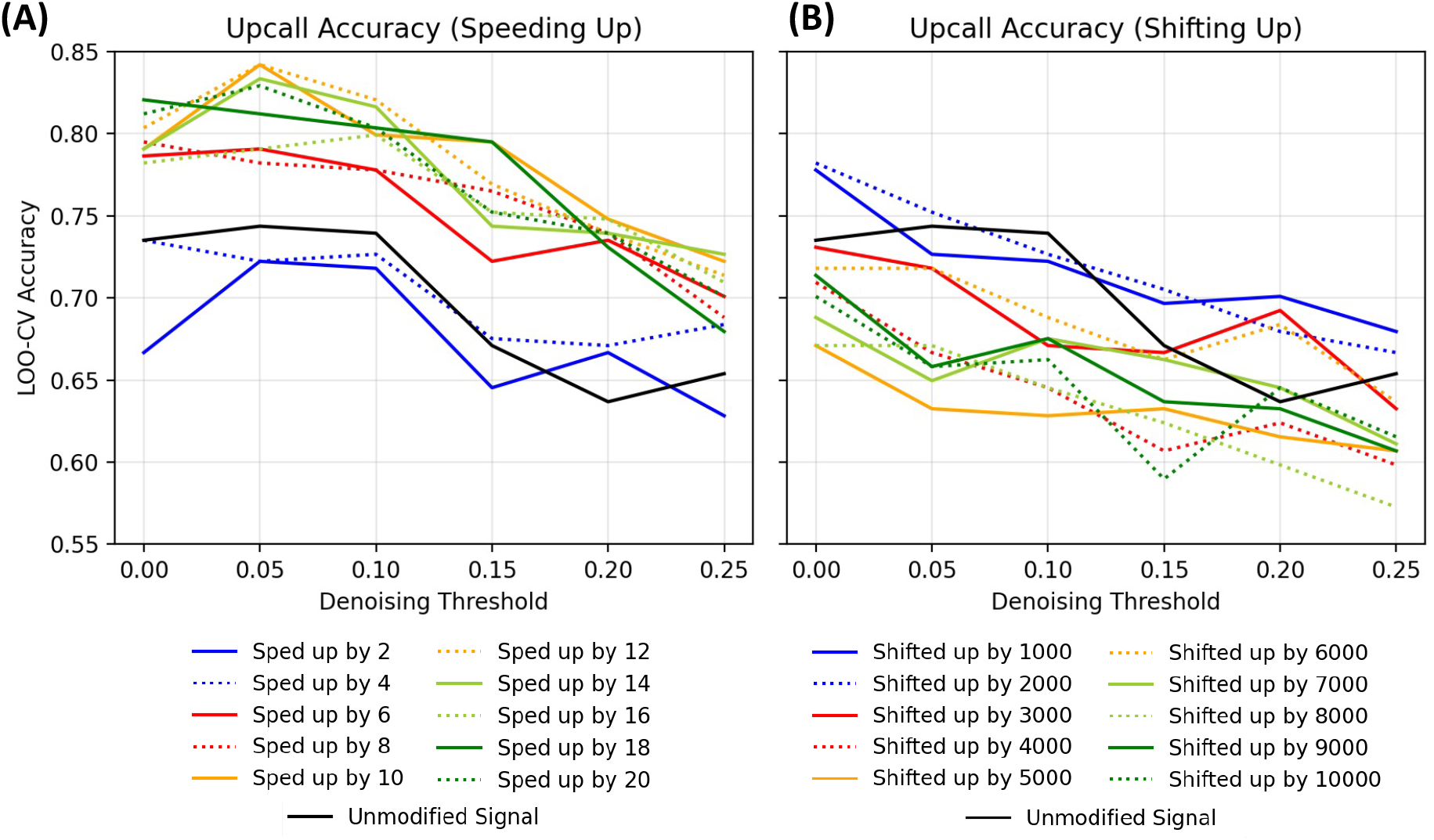
We quantified the effect of frequency modification on upcall classification accuracy, comparing speeding up the signal (**A**) and shifting up the signal with complex modulation (**B**). Overall, speeding up yields higher performance for this dataset, likely because BirdNET is more sensitive to mid-frequencies vocalized by birds.

Curiously, these results agree closely with the anticipated sensitivity of a model trained for species-level bird song and call classification. At a speedup factor of 10x, the 50-500 Hz upcall is transformed to span the 500-5000 Hz range – broadly, the range most occupied by bird calls, particularly when considering high-frequency attenuation in PAM data. Similarly, a 2 kHz shift would move the upcall to span 2050-2500 Hz. However, simply shifting up the upcall would make it appear perceptually flatter, and likely more dissimilar to the calls used for training BirdNET. On the other hand, speeding up the call would proportionately expand the frequency range of the signal, potentially emphasizing subtle frequency characteristics. Moreover, a sped-up upcall becomes structurally similar to a brief chirp or alarm call, likely creating similarities to the signals on which BirdNET was trained. Overall, these results suggest robustness of Bird-NET to new signals, while also highlighting the significance of domain shifts between training and deployment data.

### C. Classification Performance Across Thresholds

Figure 3 (left) displays the leave-one-out cross-validation accuracies of the classification pipeline as a function of the denoising threshold. Accuracies for the upcall dataset are shown in solid lines, while accuracies for the corresponding noise dataset are shown in dotted lines. Additionally, color indicates the frequency modification applied to the audio, with the unmodified spectrum shown in orange, frequency modulation by 1 kHz shown in blue, and speeding up by a factor of 10 shown in green. Overall, this figure suggests several key conclusions. When spectral subtraction is applied without thresholding (threshold = 0), we find a classification accuracy of 79% for the upcall dataset and 66% for the noise dataset. In other words, despite significant denoising, there were still sufficient tag-specific acoustic characteristics to differentiate the noise samples by tag. This finding underscores the difficulty of decoupling desired signals from correlated factors, a challenge exacerbated by the limited interpretability of deep learning methods. However, when spectral subtraction was followed by increasingly restrictive thresholding, the noise accuracy gradually decreased to the level of a random classifier, while the call accuracy remained high. Notably, at a threshold of 0.15, when the noise dataset is effectively random (24%), the call accuracy remains at around 80%, indicating that the upcall samples can be successfully differentiated by individual even after controlling for ambient noise.

**FIG. 3.**
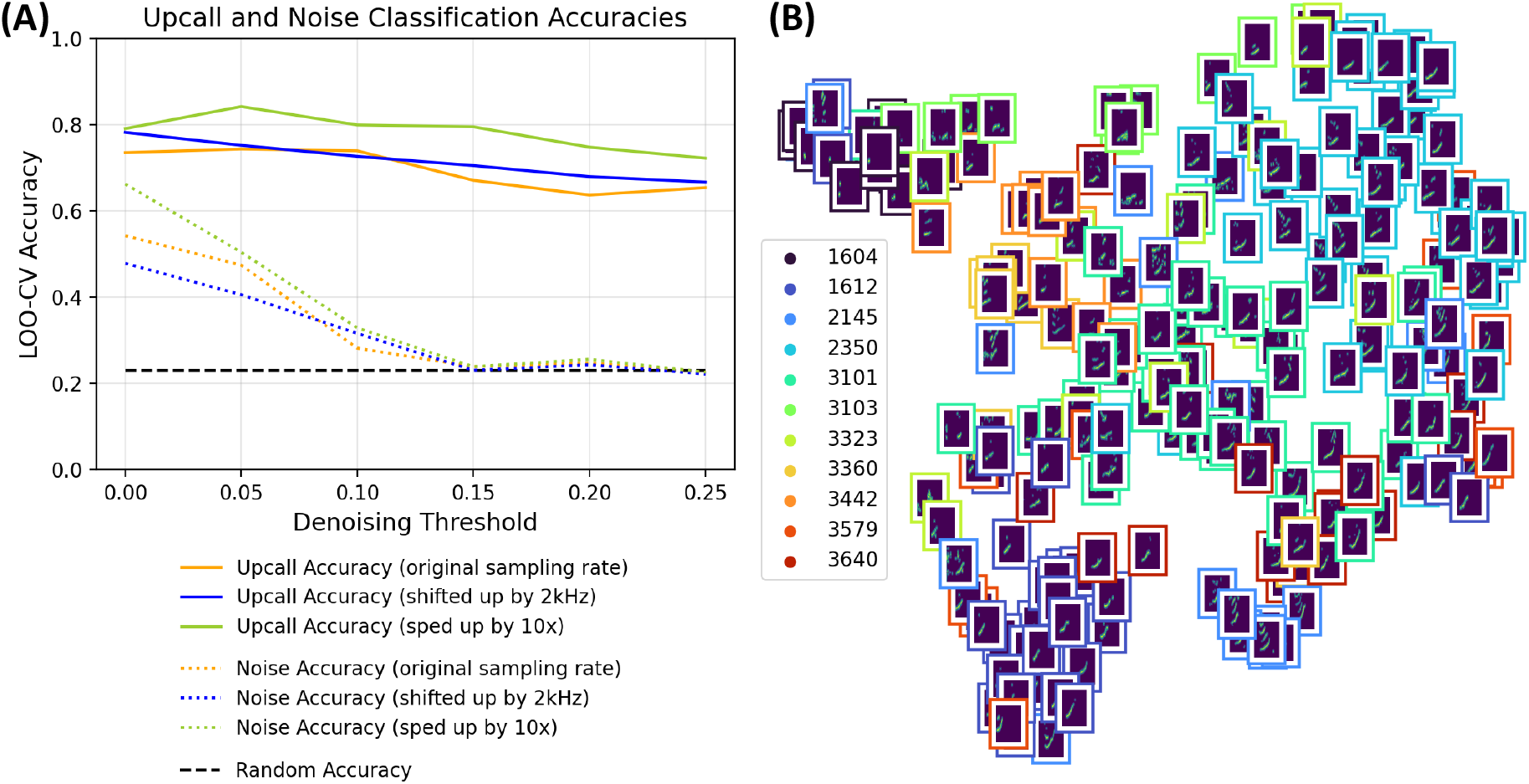
**(A)** Overall leave-one-out cross-validation accuracies for both the upcall dataset and the noise dataset at varying de-noising thresholds. **(B)** Two-dimensional UMAP projection of the upcall embeddings for a denoising threshold of 0.15. The color framing each spectrogram indicates the individual.

To visualize the distribution of intra- and inter-individual variability across upcalls, Figure 3 (right) presents a two-dimensional UMAP projection of the BirdNET embeddings for all calls at a threshold of 0.15. The colors of the spectrogram borders indicate individual identity. Overall, there is visible grouping of calls by individual whale, showing that individual variation is a major driver of variability in upcall structure. Some differences in upcall duration, slope, and frequency range are visually perceptible across the UMAP space. Finally, the smaller separate cluster at the edge of the figure is composed of very low-SNR upcall samples that lost nearly all signal structure following denoising.

### D. Effects of Ambient Noise

Figure 4 displays the accuracies for detection (in red) and recognition (in blue) as a function of SNR. At the lowest SNR (−30 dB), neither detection nor recognition is feasible. The detection curve begins to increase around −20 dB, rising sharply and starting to plateau just above −10 dB. In contrast, the recognition curve remains near-random until −15 dB, followed by a more gradual increase, reaching a plateau near 10 dB. The difference between these curves gives the difference in thresholds between detection and classification. If both curves are normalized to span the range from 0 to 1 and linearly interpolated, then this gap can be estimated at intermediate values. For instance, at the 50%, 90%, and 95% values, the SNR thresholds for detection are −15 dB, −10 dB, and −8.5 dB (respectively) while the corresponding thresholds for recognition are −7 dB, 5 dB, and 8 dB. Therefore, the threshold gaps at these values are 8 dB, 15 dB, and 16.5 dB.

**FIG. 4.**
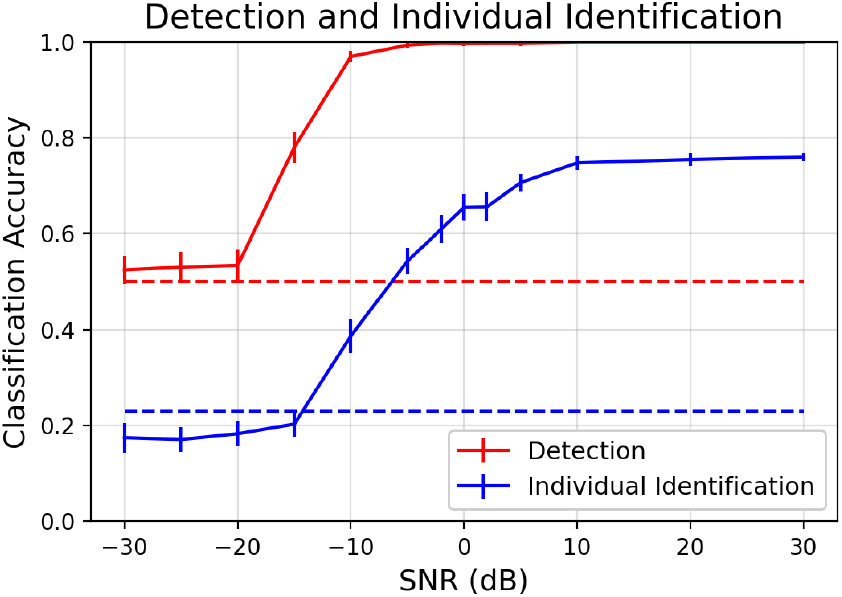
Classification accuracy of detection (in red) and individual identification (in blue) tasks across varying SNR for the upcall dataset. The solid lines indicate classification accuracies, and the dashed lines indicate random accuracy.

## V. DISCUSSION

### A. Implications for Acoustic Individual Identification

Our work contributes to, and closely aligns with, a broader context of acoustic individual identification (AIID) reviewed and conceptually formulated by Knight et al^54^. Of the 598 published studies of AIID reviewed in this work, 96% have indicated that individuals can be distinguished vocally, strongly suggesting that acoustic signatures can be found in most-sound producing taxa. However, while the review included 65 papers on “aquatic mammals”, almost all considered pinnipeds or odontecetes. Only three papers focused on individual identity in mystecetes, of which two examined humpback whales^55,56^, accompanied by the NARW analysis of McCordic et al^43^ which we continue in this work. Furthermore, of all studies considered in the review, less than 5% leveraged deep learning, likely due to the challenges of training models with low sample sizes. We hope that our work can contribute to the understanding of mysticete acoustic behavior, as well as provide a methodology for expanding the use of deep learning within AIID.

In the terminology of Knight et al, our analysis considers the “closed-set” problem of building a classifier to distinguish a known number of specific individuals, with “targeted” tag recordings that have relatively high SNR. The associated “open-set” problem — determining the number of NARW present in an area from passive acoustic data — could enable greatly streamlined abundance estimation, and therefore would be highly applicable to conservation needs. However, this task is substantially more challenging: both in acquiring a ground-truth dataset to evaluate possible techniques, and in designing an abundance estimation methodology that would be accurate and robust to irrelevant dataset characteristics. For NARW, it could be feasible to construct a ground-truth passively-recorded AIID dataset through acoustic localization, continuing prior work on abundance estimation^57^. Additionally, the denoising approach utilized here could similarly be applied to large-scale datasets to improve robustness to ambient noise conditions. However, a number of external factors may affect acoustic signatures, complicating the problem of individual identification, especially over broader spatial and temporal scales. In particular, NARW are known to refine upcall characteristics with age^43^, particularly as calves and juvenile whales mature^58^, and modify frequency and amplitude in response to ambient noise conditions^59^. Behavioral state, which in turn is closely tied to diel and seasonal cycles, could also affect upcall characteristics^33^. At different distances, effects of acoustic propagation will also attenuate and distort upcalls. Thorough understanding of these effects would be key for robust AIID-based abundance estimation.

### B. Detectability and Communication Space

The foundational study by Clark et al (2009) reformulated the SNR threshold to represent different aspects of signal recognition, terming it the “recognition differential” (RD), defined by the addition of a detection threshold (DT) term (the SNR threshold of animal auditory perception) minus a signal processing gain (SG) (the improvement in detectability and/or recognition due to signal structure) and a directivity index (DI) term (representing binaural hearing gain). This formulation is flexible, and can be adapted for different forms of communication by appropriately adjusting the latter two terms. In their analysis, the authors set DT = 10 dB, DI = 0 dB, and SG = 16 dB, yielding an overall threshold of −6 dB that defined the outer boundary of the detection space. These values were widely used in further communication space studies for a variety of baleen whales, call types, and habitats^60–62^. However, the 10 dB value was a general estimate used across sonar systems and marine mammal studies, such as from a behavioral study of a trained harbor porpoise^63^, and the 16 dB value was derived from a rule-of-thumb estimate based on the duration-bandwidth product of a signal. The wide utilization of these rough estimates underscores the data deficiency in marine mammal perception, and consequent challenges posed for assessing anthropogenic impacts.

Notably, despite the significant differences in methodology, the overall threshold of −6 dB is quite similar to the −10 dB detection threshold calculated in this work. However, our work also suggests that this threshold may not be sufficient for individual identification, which would require an additional 8-16 dB. When considered in linear units, even 8 dB (corresponding to a 50% detection probability) would carry very significant implications. In an environment in which acoustic propagation can be approximated with simple spherical spreading (10 log_10_(*r*) decrease in amplitude), an 8 dB difference in amplitude would correspond to a detection range 6.3 times greater than the individual identification range, and therefore a 2-dimensional detection space about 40 times greater than the individual ID space. Moreover, in an shallow environment in which acoustic propagation can be instead approximated with simple cylindrical spreading (5 log_10_(*r*)), an 8 dB difference in amplitude would correspond to a detection range about 40 times greater, and consequently a 2D detection space 1580 times greater than the individual ID space.

Given the critical state of the NARW population, particular attention has been dedicated to understanding the impacts of anthropogenic noise on their communication. Through a simulation grounded in real-world measurements, Clark et al (2009) found that the passage of a single commercial vessel can decrease the detection space by 97%^24^. Hatch et al (2012) analyzed passive acoustic recordings in the Stellwagen Bank National Marine Sanctuary to quantify acoustic masking due to both heightened ambient noise and shipping noise across a range of parameters^61^. Broadly, the authors find that relative to hypothesized historical ambient noise levels, the heightened underwater noise levels have led to 65% loss of the detection space during quiet periods, and 87% loss during noisy periods. By modeling acoustic propagation, Tennessen and Parks (2016) calculated that right whale upcalls will be entirely masked by noise if a container ship is within a 25 km range^41^. While right whales could compensate by increasing their call amplitude by 10 or 20 dB, this translates to a ten-fold or hundred-fold increase in amplitude: likely a significant energetic cost. Crucially, all of these studies ultimately evaluate the ability to detect calls, and not the capacity to perceive vital, more complex information such as individual identity. Computational estimation of communication parameters could begin to address these unknowns, and refine assessments of anthropogenic impact.

### C. Challenges and Caveats

Our analysis leverages the properties of audio embeddings as a proxy for the perceptive capabilities of NARW. While this is clearly an imperfect comparison, recent studies evaluating the relative capabilities of deep neural networks (DNN) and biological auditory perception suggest strong parallels between the two. In broader research on audio processing, DNNs have been shown to match and even outperform human participants across diverse audio classification tasks. When compared against traditional models of the auditory cortex, DNNs have been shown to better predict human auditory response^64^. Moreover, activations across a variety of DNNs show correspondences to different brain regions indicated by functional Magnetic Resonance Imaging (fMRI) scans, suggesting that deep networks consistently learn similar pathways of underlying computational processing^64,65^. Given the lack of data on auditory perception in wild animals, and the difficulty of acquiring it, deep learning could be beneficial for estimating quantitative parameters in the study of animal communication.

In this work, we quantified information transfer in the form of individual identification from a contact call. Of course, information is a highly context-dependent concept, and will vary by the caller, listener, signal characteristics, and form of information conveyed. Additionally, these results represent SNR in the presence of homogenous white noise; in practice, heterogeneous noise characteristics would otherwise affect these results. Lastly, SNR may have variable definitions, so quantitative values may not directly transfer across studies. Yet even with these simplifications, the resulting detection threshold closely matched values from prior studies.

Deep learning opens new opportunities for bioacoustic analysis, but not without challenges and caveats. In this study, tag-specific ambient noise proved to be a significant confounding factor. When applying highly sensitive machine-learning-based tools, caution is warranted, particularly when ecological conclusions may be impacted by correlated patterns of ambient noise. Additionally, pre-trained models can effectively analyze small-scale datasets, enabling classification performance that would have been infeasible with prior machine learning methods. However, domain differences between the training and testing datasets may affect performance. In this study, we confirmed that a model trained on bird call and song recognition could be successfully applied to low-frequency whale calls, but found that scaling the calls to a bird-like frequency range improved classification accuracy.

## VI. CONCLUSION

In this study, we demonstrate that deep-learning-based techniques can distinguish individual identity in a dataset of NARW upcalls – a call type considered to be heavily stereotyped. Moreover, by leveraging deep learning, we quantify the ability to recognize individual characteristics in a noisy environment, and thereby infer the loss of NARW communication space due to increased ocean noise levels. Overall, while our analysis focused on one species of critical conservation concern, the methods are general, and can be applied across taxa. In particular, we hope that this study can serve as an example of a low-cost method for quantifying communication parameters for a given species and/or call type, thereby improving estimates of acoustic masking and corresponding anthropogenic impact assessments.

